# Tropism of SARS-CoV-2 for Developing Human Cortical Astrocytes

**DOI:** 10.1101/2021.01.17.427024

**Authors:** Madeline G. Andrews, Tanzila Mukhtar, Ugomma C. Eze, Camille R. Simoneau, Yonatan Perez, Mohammed A. Mostajo-Radji, Shaohui Wang, Dmitry Velmeshev, Jahan Salma, G. Renuka Kumar, Alex A. Pollen, Elizabeth E. Crouch, Melanie Ott, Arnold R. Kriegstein

**Author notes:** These authors contributed equally (Madeline G. Andrews, Tanzila Mukhtar).

## Abstract

The severe acute respiratory syndrome coronavirus 2 (SARS-CoV-2) readily infects a variety of cell types impacting the function of vital organ systems, with particularly severe impact on respiratory function. It proves fatal for one percent of those infected. Neurological symptoms, which range in severity, accompany a significant proportion of COVID-19 cases, indicating a potential vulnerability of neural cell types. To assess whether human cortical cells can be directly infected by SARS-CoV-2, we utilized primary human cortical tissue and stem cell-derived cortical organoids. We find significant and predominant infection in cortical astrocytes in both primary and organoid cultures, with minimal infection of other cortical populations. Infected astrocytes had a corresponding increase in reactivity characteristics, growth factor signaling, and cellular stress. Although human cortical cells, including astrocytes, have minimal ACE2 expression, we find high levels of alternative coronavirus receptors in infected astrocytes, including DPP4 and CD147. Inhibition of DPP4 reduced infection and decreased expression of the cell stress marker, ARCN1. We find tropism of SARS-CoV-2 for human astrocytes mediated by DPP4, resulting in reactive gliosis-type injury.

## Introduction

The novel coronavirus, SARS-CoV-2, causes the life-threatening illness, COVID-19, and is responsible for the current global pandemic resulting in over 1.5 million deaths world-wide. Although SARS-CoV-2 infection most notably impairs respiratory function, the capacity to infect other cell types and disrupt function of other vital organ systems is currently being elucidated. Strikingly, many patients suffering with or having recovered from COVID-19 present a range of neurological symptoms from seizures, encephalopathy, stroke, headaches, dizziness, short-term memory loss, loss of smell and taste and complaints of mental sluggishness or inability to focus (*1, 2*). Individuals with mental health diagnoses are also more susceptible to coronavirus infection and have impaired long-term health outcomes (*3*).

It is unclear whether this range of neurological symptoms is a result of direct infection of the neural tissue, or a secondary consequence of widespread inflammation downstream of viral infection in other tissues. Strokes, for example, result from SARS-CoV-2 induction of prothrombotic autoantibodies and a resulting clotting disorder. But SARS-CoV-2 may also be able to directly infect cells in the brain. SARS-CoV-2 can infect neurons in the nasal epithelium, a potential mode of entry to the CNS from the periphery, and presence of viral RNA has been detected in neural tissues in patients (*4*). Recent studies report mixed findings concerning the presence of coronavirus viral RNA and antibodies in the cerebral spinal fluid (CSF) of COVID-19 patients (*5, 6*). However, choroid plexus organoids containing the cell type that produces CSF, can readily be infected by SARS-CoV-2 *in vitro* (*7, 8*). Together, these studies suggest the capacity for viral transmission into the CNS through the nasal epithelium and/or CSF. Additionally, the virus could potentially infect and disrupt the brain vasculature. Studies of postmortem brain tissue from COVID-19 patients have observed widespread inflammation in the brainstem characterized by infiltration of immune cells including microglia and T cells, as well as infection of cranial nerves, microvascular injury, fibrinogen leakage, and extensive astrogliosis (*9, 10*). Moreover, viral RNA has recently been observed in cortical astrocytes from post-mortem patient tissue (*11*).

The impact of SARS-CoV-2 infection on the developing brain is uncertain. While pediatric symptoms of COVID-19 are typically milder than in adults, infants have increased risk for infection and severity of adverse complications than older pediatric patients, with as many as 30% of infected newborns requiring intensive care (*12–16*). Almost one-fifth of all reported COVID-19 pediatric cases are infants (*17*) and many neonates with infected mothers also test positive for SARS-CoV-2 within the first few days of life (*18*). An outstanding question remains regarding the transmission route for SARS-CoV-2 in newborns and whether infection can occur *in utero*. While the majority of amniotic fluid, cord blood, and placental tissue samples test negative for SARS-CoV-2 RNA (*19*), several case studies have reported maternal to fetal transmission during gestation with high levels of infection in the placenta (*20–22*). In one particular case, the newborn demonstrated significant neurological consequences including gliosis in cortical and subcortical structures (*20*).

The vulnerability of particular cell types in the developing brain and the impact on brain health and function require further study and human stem cell-derived neural models are a tractable method to evaluate viral tropism (*23*). Data from cerebral organoids, which are reflective of developmental stages, suggest that *in vitro* neurons may be vulnerable to SARS-CoV-2 infection. However the results regarding susceptibility of neurons of different brain regional identities have been mixed across studies and the capacity to infect other neural cell types, including glia, has not been extensively explored (*7, 8, 24*). Here we utilized primary developing human cortical tissue and cortical organoid models across neurogenic and gliogenic stages to evaluate whether human neural cells can be infected by SARS-CoV-2. In cortical tissue cultures and cortical organoids exposed to SARS-CoV-2 we find significant infection and viral replication in astrocytes, but minimal infection in other neural cell types. Developing cortical tissue expresses very little ACE2 receptor, suggesting that the virus may use another means of viral entry such as SARS-CoV-2 co-factors, DPP4 or CD147, which we find highly expressed in infected astrocytes. Inhibiting DPP4 reduces infection, suggesting DPP4 is a mode of entry into developing astrocytes. Our study provides evidence of tropism for specific human neural cell types with implications for the vulnerability of the developing brain and the potential for neural infection after SARS-CoV-2 exposure in postnatal life.

## Results

To evaluate the capacity of SARS-CoV-2 to directly infect human cortical cells, we exposed organotypic slice cultures of developing human cortical samples to SARS-CoV-2 for two hours and collected cultures 72 hours later (Figure 1A). Infected cells were identified with a coronavirus nucleocapsid (N) antibody and viral replication by the presence of double stranded RNA (dsRNA). 100% of infected cells were co-labeled, indicating both infection and replication (Figure 1B). To evaluate the vulnerability of neural cell types for infection we utilized markers for cortical progenitors, neurons, and glial cells, including astrocytes and oligodendrocytes in infected cultures. Strikingly, we found abundant infection in astrocytes throughout the outer subventricular zone (oSVZ), as indicated by the presence of viral N protein and dsRNA in cells expressing glial fibrillary acidic protein (GFAP) and Aquaporin 4 (AQP4) (Figure 1B). However, we observed minimal infection of other cell types present in the developing cortex, in particular, NEUN+ neurons and KI67+ dividing cells were rarely infected (Figure 1C). Immature astrocytes expressing PAX6 and HOPX, as well as mature astrocytes expressing S100B, showed high infectability (S Figure 1A). Other glial cell types, including OLIG2+ oligodendrocyte precursor cells (OPC) and the resident neural immune cells, IBA1+ microglia, had little infection (S Figure 1B). Cells of the cortical excitatory neural lineage, including neurogenic radial glia and intermediate progenitor cells (IPC), demonstrated a lack of infection, confirming an infection bias toward glial over neuronal cell types or neuronal progenitors (S Figure 1C).

**Figure 1.**
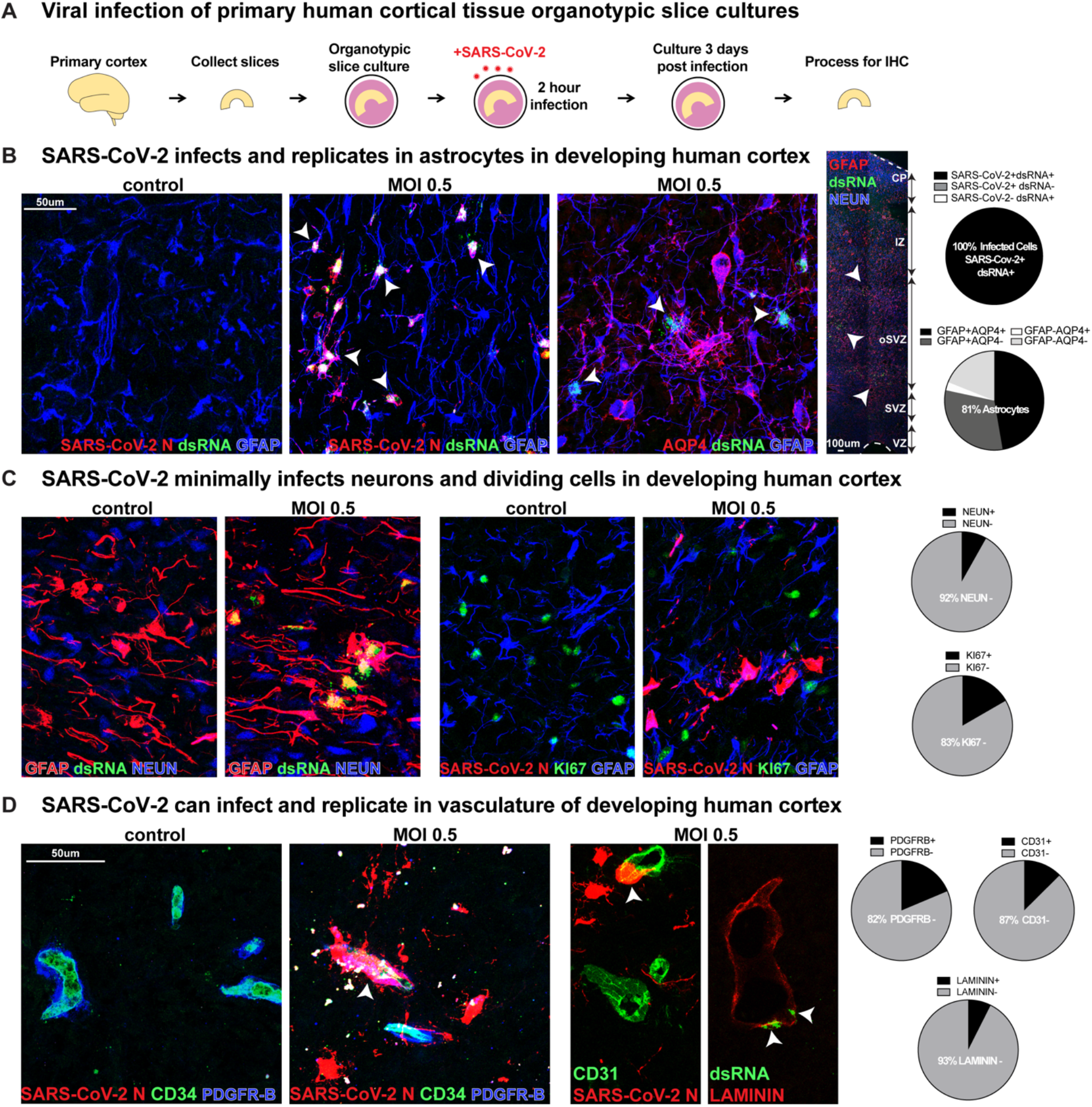
SARS-CoV-2 Infects Astrocytes in Developing Human Cortex. **A)** Experimental paradigm for viral infection of human cortical tissue. Primary cortical tissue was acutely sectioned and cultured at the air-liquid interface. The next day tissue was infected with SARS-CoV-2 at MOI 0.5 and cultured for 72 hours before being collected and processed. **B)** SARS-CoV-2 infects GFAP+AQP4+ astrocyte cells in the developing human cortex, which are predominantly located in the subventricular zone (SVZ), where >81% of cells assayed expressed markers of astrocytes. 100% of infected cells expressed both SARS-CoV-2+ nucleocapsid (N) antibody and double-stranded (ds)RNA antibody. White arrowheads indicate dsRNA+GFAP+ infected astrocytes (dsRNA+SARS-CoV-2 N+ n=31 cells, GFAP+AQP4+ n=74 cells across two biological samples and four technical replicates). **C)** Few other neural types were infected, as indicated by co-labeling of SARS-CoV-2 N or dsRNA, where <8% of cells assayed expressed NEUN, a marker of cortical neurons, and <17% expressed KI67, a marker of dividing cells (NEUN n=613 cells, KI67 n=186 cells across two biological samples and four technical replicates). **D)** Vascular cell types can also be infected where 7% of LAMININ+ blood vessels, 13% of CD31 + endothelial cells and 18% of PDGFR-β+ mural cells have infection. White arrowheads indicate infected vascular cells (Laminin n=269 cells, CD31 n=247 cells, PDGFRB=225 cells across two biological samples and four technical replicates).

To evaluate a potential route of viral entry to the brain, we explored infection in vascular cell types. Developing blood vessels, which express Laminin, are composed of an inner layer of endothelial cells (expressing CD31 and CD34) and an exterior layer of mural cells (expressing PDGFR-β) (Figure 1D). Approximately 13% of infected cells (dsRNA+ or N+) were CD31+ endothelial cells and 18% were PDGFR-β+ mural cells, indicating a capacity for SARS-CoV-2 infection in vascular cell types. Although the proportion of infected vascular cells is much lower than cortical astrocytes, the infected vasculature consisted of small blood vessels located in the SVZ. Cells adjacent to infected mural and endothelial vascular cells were also frequently infected, suggesting a potential means of viral spread to cortical astrocytes, which are key in blood-brain barrier formation.

To evaluate the capacity for infection over a range of developmental stages and to further explore viral tropism, we utilized cortical organoids infected with SARS-CoV-2 at stages of early neurogenesis (week 5), peak neurogenesis (week 10), early gliogenesis (week 16) and peak gliogenesis (week 22) (Figure 2A). We observed that organoids from neurogenic and early gliogenic stages were rarely infected, and the infrequently infected cells did not express markers of SOX2+ cortical progenitors, NEUN+ neurons or glial lineage markers, suggesting an off-target or non-cortical lineage (Figure 2B). At week 16, we observed infection of a few GFAP+ cells in one stem cell line, but infection was absent in a second stem cell line at this time point. Overall, at neurogenic stages in organoids, we found rare viral N+, but not dsRNA+, cells, indicating low levels of entry without evidence of further replication or productive infection (Figure 2C). However, by week 22 of differentiation, a stage where significant gliogenesis has occurred, GFAP+AQP4+ astrocytes were robustly infected, consistent with our results in primary tissue samples. NEUN+ cortical neurons were rarely infected (Figure 2D-E). Together, the primary and organoid cortical findings suggest SARS-CoV-2 can readily infect astrocytes in the developing human cortex.

**Figure 2.**
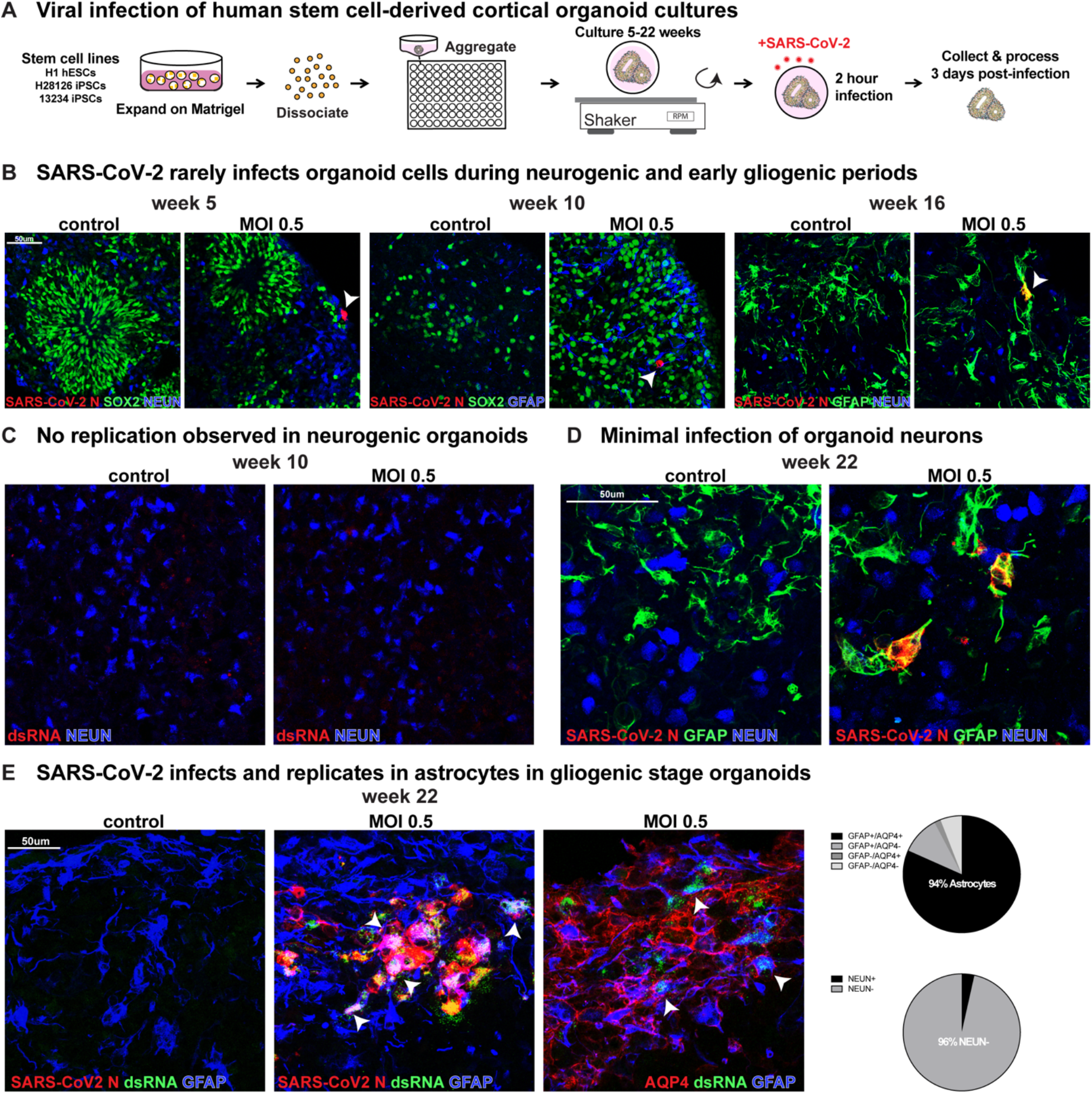
SARS-CoV-2 Infects Astrocytes in Cortical Organoids. **A)** Viral infection of cortical organoids. Human stem cells from several lines were aggregated and differentiated toward cortical identity in suspension. After 5, 10, 16 or 22 weeks of differentiation, organoids were exposed to SARS-CoV-2 for 2 hours, media was replaced and then collected 72 hours later. **B)** After 5, 10, or 16 weeks organoids only indicated rare infection (white arrowheads). At five and 10 weeks, SARS-CoV-2 N+ cells did not co-express SOX2, NEUN or GFAP indicating infected cells are not cortical progenitors, neurons or astrocytes and may instead be an off-target population. However, after 16 weeks, in one stem cell line a few GFAP+ astrocytes were infected. **C)** Although rare cells are infected at neurogenic stages, as indicated by coronavirus N antibody presence, there was no observed viral replication with dsRNA at these timepoints. **D)** At late gliogenic stages, by week 22 infection was readily observed in GFAP astrocytes but not NEUN+ neurons. **E)** 94% of infected cells stained positive for markers of astrocytes GFAP or AQP4, while only 4% are NEUN+ neurons. White arrowheads indicate SARS-CoV-2+ dsRNA+ GFAP+ AQP4+ astrocytes (GFAP+AQP4+ n=169 cells, NEUN n=143 cells).

To assess the functional consequences of SARS-CoV-2 infection on developing astrocytes, we assayed the abundance of the cell death marker, Cleaved Caspase-3. We observed no increase in apoptosis in primary cortical cultures 72 hours post-infection in dsRNA+ GFAP+ cells (Figure 3A). In a recent study of adult post-mortem samples, infected astrocytes were observed to have impaired protein folding, translation initiation, and metabolic function (*11*). We investigated the impact of SARS-CoV-2 on cellular stress by evaluating the abundance of endoplasmic reticulum (ER) stress and the unfolded protein response (UPR) gene, ARCN1. ARCN1 is lowly expressed in developing human cortex tissue, but after infection by SARS-CoV-2, we found abundant expression in infected glial cells (Figure 3B).

**Figure 3.**
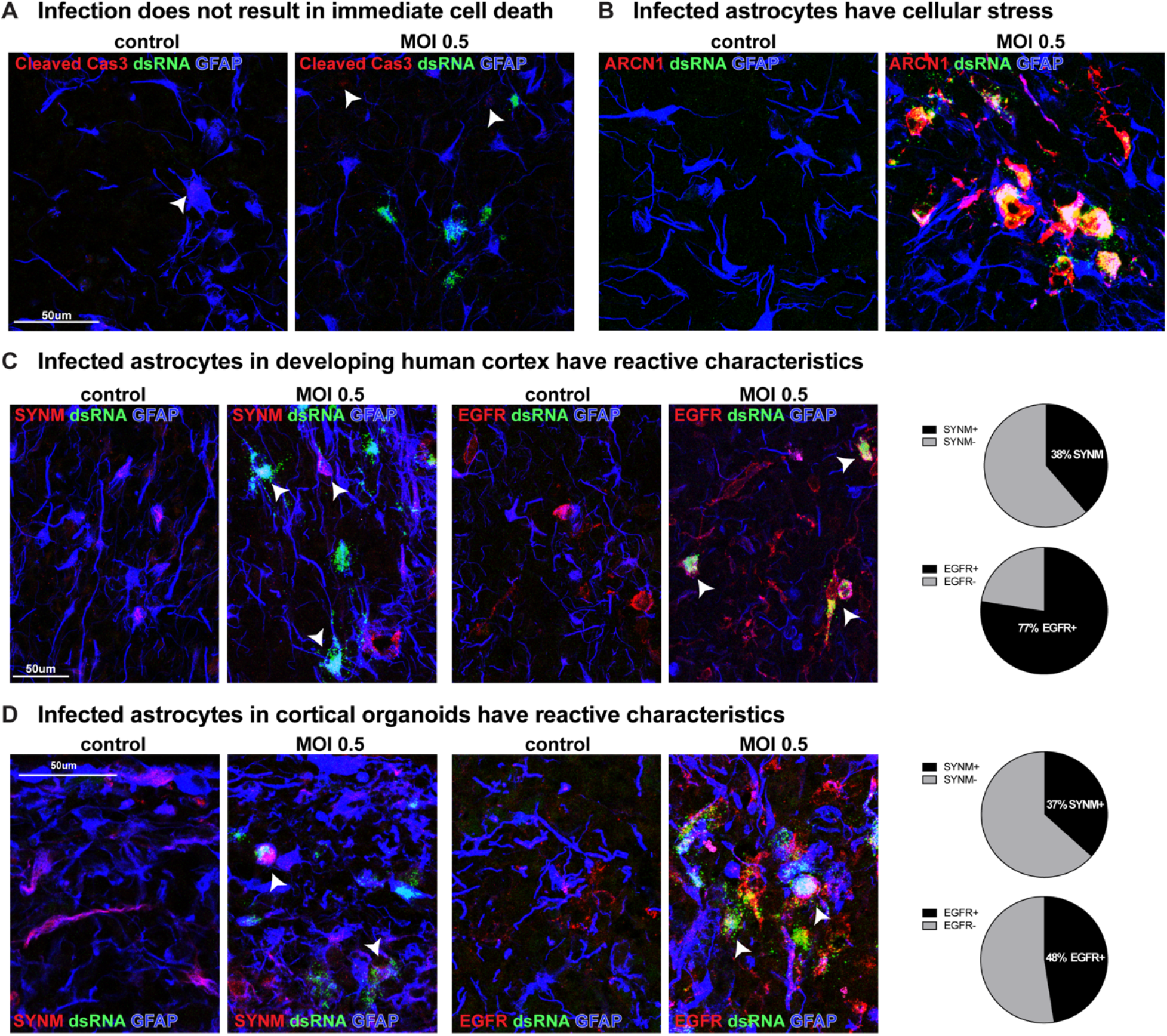
SARS-CoV-2 Infection Increases Cell Stress and Reactivity in Cortical Astrocytes. **A)** After SAR-CoV-2 infection, there is no immediate increase in cell death in infected cells, as indicated by Cleaved Caspase 3 staining. **B)** Infected cells have an increase in cell stress, as indicated by the ER stress marker, ARCN1. **C)** Approximately one-third of infected astrocytes in primary cortical tissue express the reactive marker SYNM and three-quarters have the receptor for growth factor signaling, EGFR (SYNM n=62 cells, EGFR n=142 cells). **D)** The same proportion of infected organoid cells express SYNM and about one-half express EGFR (SYNM n=172 cells, EGFR n=143 cells).

We sought to evaluate whether infected astrocytes have atypical characteristics, including increased reactivity or activated growth factor signaling, common after SARS-CoV-2 infection (*25*). Astrocytes can express the reactive marker, Synemin (SYNM), under control conditions, but more than 1/3 of infected astrocytes expressed SYNM, indicating either preferential infection by SARS-CoV-2 of reactive astrocytes or induction of a reactive-like state post-infection (Figure 3C). Additionally, because inhibition of growth factor receptors can prevent SARS-CoV-2 replication (*26*), we evaluated the presence of Epidermal growth factor receptor (EGFR) on infected astrocytes. EGFR is involved in the regulation of inflammatory states and is expressed by some glial precursor cell types. We observed that more than 75% of infected cells expressed EGFR, and overall abundance of EGFR increased after SARS-CoV-2 infection compared to control conditions (Figure 3C). Organoid-derived astrocytes also exhibited reactive characteristics in response to infection in similar proportions to primary astrocytes, with approximately one-third of organoid-derived infected cells expressing SYNM and half expressing EGFR (Figure 3D).

The robust capacity for infection of cortical astrocytes across model systems highlights questions of viral tropism. The ACE2 receptor, the key entry factor for coronaviruses including SARS-CoV2, has low expression in the developing human cortex. Re-analysis of our published single cell RNA sequencing (sc-RNA-seq) dataset of primary human cortical development (*27*) demonstrated essentially null expression in any cell type in the developing human cortex (S Figure 2A) (bit.ly/cortexSingleCell). We further examined expression of ACE2 in bulk RNA samples from developing human cortex, adult human cortex, cortical organoids, and developing lung tissue, and observed very little expression in neural tissue compared to lung (S Figure 2A).

To determine whether there is a divergence between ACE2 RNA expression and protein abundance in human cortical cells, we immunolabeled infected cortical tissue and organoids. No ACE2 was detected across cortical samples or in dsRNA+ infected cells, when compared to ACE2-expressing positive control cells (Figure 4A). NRP1 has also been demonstrated to be a host factor for SARS-CoV-2 infection (*28*). We observed RNA expression specifically in cortical neurons, which, however, is a population where we observe minimal infection. When we evaluated NRP1 protein abundance in tissue, we similarly observed membrane-bound staining in neurons in the cortical plate (CP), but none in dsRNA+ infected cells in the SVZ (Figure 4A, S Figure 2A).

**Figure 4.**
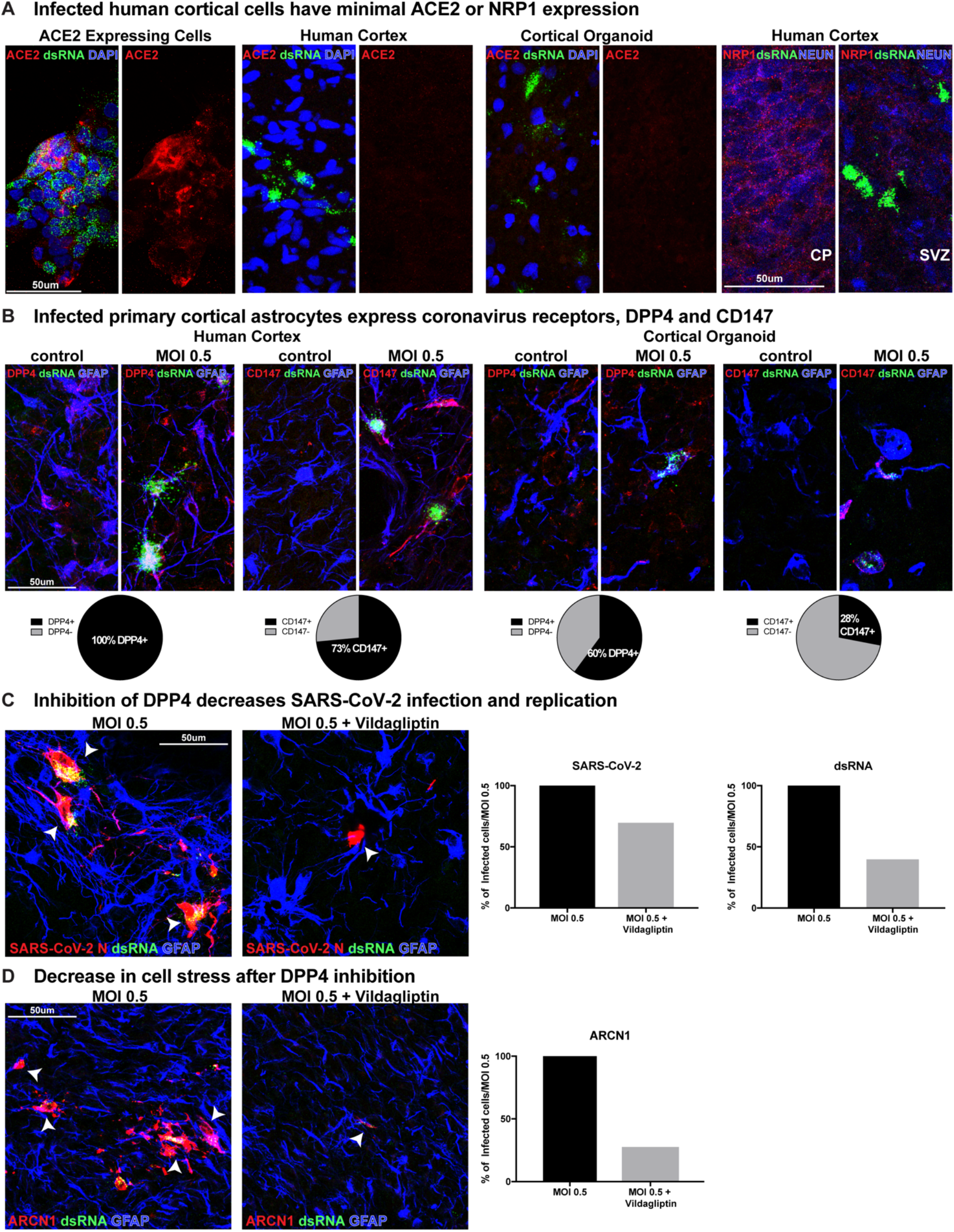
Coronavirus Receptors, DPP4 and CD147, but not ACE2 are Expressed in Developing Human Cortex. **A)** ACE2 expressing cells are readily infected by SARS-CoV-2, where they have abundant ACE2 and dsRNA presence. Primary cortical tissue and cortical organoids do not have observable ACE2 protein in cortical tissue or infected cells. NRP1 is present in cortical neurons, but not in infected cells. **B)** Infected cortical astrocytes in the SVZ of primary cortex abundantly express coronavirus receptors DPP4 and CD147, where 100% of cells assayed have DPP4 and 73% have CD147. In cortical organoids, 60% are DPP4+ and 28% are CD147+ (Primary: DPP4 n=61cells, CD147 n=83 cells; Organoid: DPP4 n=296 cells, CD147 n=239 cells) **C)** Inhibition of DPP4 by Vildagliptin results in a 30% decrease in the number of SARS-CoV-2 N+ cells and a 70% reduction in dsRNA+ cells. SARS-CoV-2 N+ dsRNA+ cells are indicated by white arrowheads (SARS-CoV-2 MOI 0.5 n=1273 cells, MOI 0.5+Vildagliptin n=879 cells, dsRNA MOI 0.5 n=571 cells, MOI 0.5+Vildagliptin n=227 cells across two biological samples and three technical replicates). **D)** ARCN1 is broadly expressed in SARS-CoV-2 infected samples, which is reduced by 70% after DPP4 inhibition by Vildagliptin. ARCN1+ dsRNA+ cells indicated by white arrowheads (MOI 0.5 n=1224 cells, MOI 0.5+ Vildagliptin n=331 cells across two biological samples and three technical replicates).

Recent studies have demonstrated that co-mediating factors are involved in SARS-CoV-2 infection. Expression of specific receptors, restriction enzymes, and proteases across tissues may be vital for mediation of SARS-CoV-2 infection in collaboration with, or perhaps independent of, ACE2 (*28–31*). We evaluated whether the extracellular glycoproteins and coronavirus receptors, DPP4 and BSG/CD147, are expressed in our primary developing human sc-RNA-seq dataset and observed broad expression in a variety of cell types and high bulk RNA expression across cortical organoids, developing human cortex, and adult cortex samples (S Figure 2A). Additionally, coronavirus restriction factors, such as LY6E and IFITM1, and proteases, TMPRSS5, FURIN, CTSB, have differential expression in cortical cell types across tissue samples and may also mediate infection (S Figure 2B, S Figure 3).

Given the absence of ACE2 or NRP1 expression in cortical astrocytes, we investigated the abundance of the coronavirus receptors, DPP4 and CD147 in our infected primary tissue samples. We observed 100% co-labeling of DPP4 and 73% of CD147 in dsRNA+ infected cells (Figure 4B). Although cortical organoids did not have the same high abundance of receptor expression as primary samples, infected organoid cells also demonstrated broad co-expression of DPP4 and CD147 (Figure 4B).

Due to the abundance of DPP4 in astrocytes prior to infection, and recent findings suggesting BSG/CD147 may not directly bind SARS-CoV-2 (*32*), we focused our attention on whether DPP4 could mediate cortical astrocyte tropism. Organotypic slice cultures were treated with the DPP4 inhibitor, Vildagliptin, for 24 hours prior to and throughout the duration of infection. There was robust infection of GFAP+ astrocytes in virally-exposed samples after 72 hours, however there was a 30% reduction in the number of SARS-CoV-2 N+ and a 60% reduction in dsRNA+ cells in the tissue samples treated with Vildagliptin (Figure 4C). Additionally, after DPP4 inhibition, there was a 70% reduction in the number of cells expressing ARCN1, indicating a decrease in activation of cellular stress (Figure 4D). Together, these data suggest SARS-CoV-2 tropism for astrocytes in the developing human cortex. Furthermore, infection increases cellular stress and reactivity, and the capacity for infection is mediated, in part, by the DPP4 extracellular receptor.

## Discussion

Our study exploring neural tropism of the novel coronavirus, SARS-CoV-2, shows that human cortical astrocytes can be directly infected and display downstream cellular stress and glial reactivity. We additionally find that vascular cell types, including endothelial and mural cells, can be infected. Vascular cells are positioned adjacent to astrocytes in blood-brain barrier formation and their close proximity provides a potential mode of viral entry into the central nervous system.

Although SARS-CoV-2 infection of airway epithelia and lung parenchyma is predominantly mediated by the ACE2 receptor, this study and others, provide evidence that alternative coronavirus receptors can also mediate infection (*28*). The capacity of SARS-CoV-2 to infect cell types that express low levels of ACE2 by utilizing other extracellular glycoproteins, suggest a broad capacity to infect a variety of cell types using a range of receptors across many organ systems, ultimately inflicting severe, systemic damage. Our findings suggest that DPP4 is a key receptor mediating infection in cortical astrocytes. However, DPP4 is not the sole receptor mediating infection, as infection is not completely eliminated after DPP4 inhibition. Though we did not observe high expression of canonical proteases TMPRSS2/4 in human cortex, other proteases TMPRSS5, FURIN and CTSB are widely expressed, suggesting potential alternate modes of viral entry in the brain. Exploration for additional novel SARS-CoV-2 receptors and proteases will help determine the range of entry factors utilized by the virus and how it may impact particular cellular populations.

We observed a strong tropism of the SARS-CoV-2 virus for astrocytes, a key ‘support cell’ that regulates a myriad of vital functions in the developing and adult brain. Astrocytes regulate neurotransmitter concentration and re-uptake for appropriate synaptic communication, they mediate blood-brain barrier function, and regulate neural metabolism and inflammation (*33*). Disorders of astrocyte development can have devastating impacts on brain function. For example, astrocyte dysfunction in Alexander Disease results in increased cellular stress and autophagy, changes to astrocyte morphology, inability to regulate the concentration of neurotransmitters and ions for appropriate synaptic activity, and decoupling of astrocytes to one another, blood vessels, and the neuronal microenvironment (*33*). These cellular changes result in seizures, inability to control motor function, and ultimately death. The devastating consequences of Alexander Disease highlight the vital importance of appropriate astrocyte development to regulate basic synaptic function and overall brain activity and health.

ACE2 and the protease, TMPRSS2, are highly expressed in the trophectoderm of the preimplantation embryo, which develops into the placenta. Trophoblast cells also express the receptors DPP4 and CD147 and low levels of the restriction factors IFITM and LY6E, rendering placental cells permissible to viral entry (*30*). Trophectoderm cells can be infected *in vitro*, supporting the potential for vertical transmission of SARS-CoV-2 from infected mother to fetus.

Case studies of placental infection and newborns who test positive for SARS-CoV-2 validate the capacity for viral transmission across the maternal-fetal interface and access to developing organ systems (*22*). However, the long-term impact of gestational infection during early development is yet to be determined. Future studies should explore mechanisms of viral transmission to different cell types and structures in the human brain and how vulnerable different neural cells are to SARS-CoV-2 infection across different stages of life.

Our findings suggest that developing astrocytes are particularly vulnerable to SARS-CoV-2, however another recent study indicates that adult astrocytes may also be susceptible to infection and that other coronavirus receptors, like CD147, are abundant in mature astrocytes (*11, 30*). We observe an increase in CD147 expression upon infection, suggesting a potential positive feedback loop, as observed in case of ACE2 in lung alveolosphere infection (*34*). Glial vulnerability to SARS-CoV-2 may extend across the lifetime but can be particularly consequential at early stages of brain development. While vertical transmission between COVID+ mothers and their neonates may be an infrequent event, there is a paucity of literature studying the sequelae of severe COVID cases in pregnancy (*35*). It will be critical to determine the long-term prevalence of gliosis as well as the incidence of neurological disease, particularly seizures, in infected neonates.

In this study, we performed a comprehensive analysis of primary tissue, in parallel with organoid models at a range of developmental timepoints. While cerebral organoid models have limitations in terms of the diversity and fidelity of neural cell types (*36*), our studies indicate that astrocytes are preferentially infected in both primary human cortical tissue and cortical organoid models. Moreover, organoid-derived and primary astrocytes had similar proportions of receptor abundance and reactivity phenotypes following infection, further emphasizing the utility of organoid models to study viral tropism and cellular responses. However, future studies exploring post-mortem patient neural tissues will be required to understand if mature astrocytes are as vulnerable as immature astrocytes, and to further define the receptors that may mediate infectivity. The results of our qPCR analysis indeed suggest moderate expression of coronavirus receptors ACE2, DPP4, CD147, and NRP1 in the adult cortex. The range of COVID-19 associated neurological symptoms including dizziness, seizures, and cognitive difficulties, may reflect the involvement of astrocytes that are vital to global brain homeostasis and function. Further studies elucidating potential routes of CNS infection as well as strategies to block viral entry may aid in controlling transmission and help alleviate disease symptoms.

## Acknowledgements

The authors thank Qiuli Bi, William Walantus, Maureen Galvez, Jayden Ross, Mohini Bade, Ethan Winkler, Mercedes F Paredes, Tomasz J Nowakowski, Faranak Fattahi, Jonathan Ramirez, Eric Huang, and members of the ARK laboratory for providing resources, technical help, and useful discussions. This study was supported by NIH awards U01MH114825 and R35NS097305 to ARK, 5DP1DA038043 to MO, and K99MH125329 to MGA, the Brain & Behavior Research Foundation NARSAD Young Investigators Award to MGA and the Swiss National Science Foundation Early Postdoc Mobility fellowship to TM. MO acknowledges support through a gift from the Roddenberry Foundation.

---

**S Figure 1.**
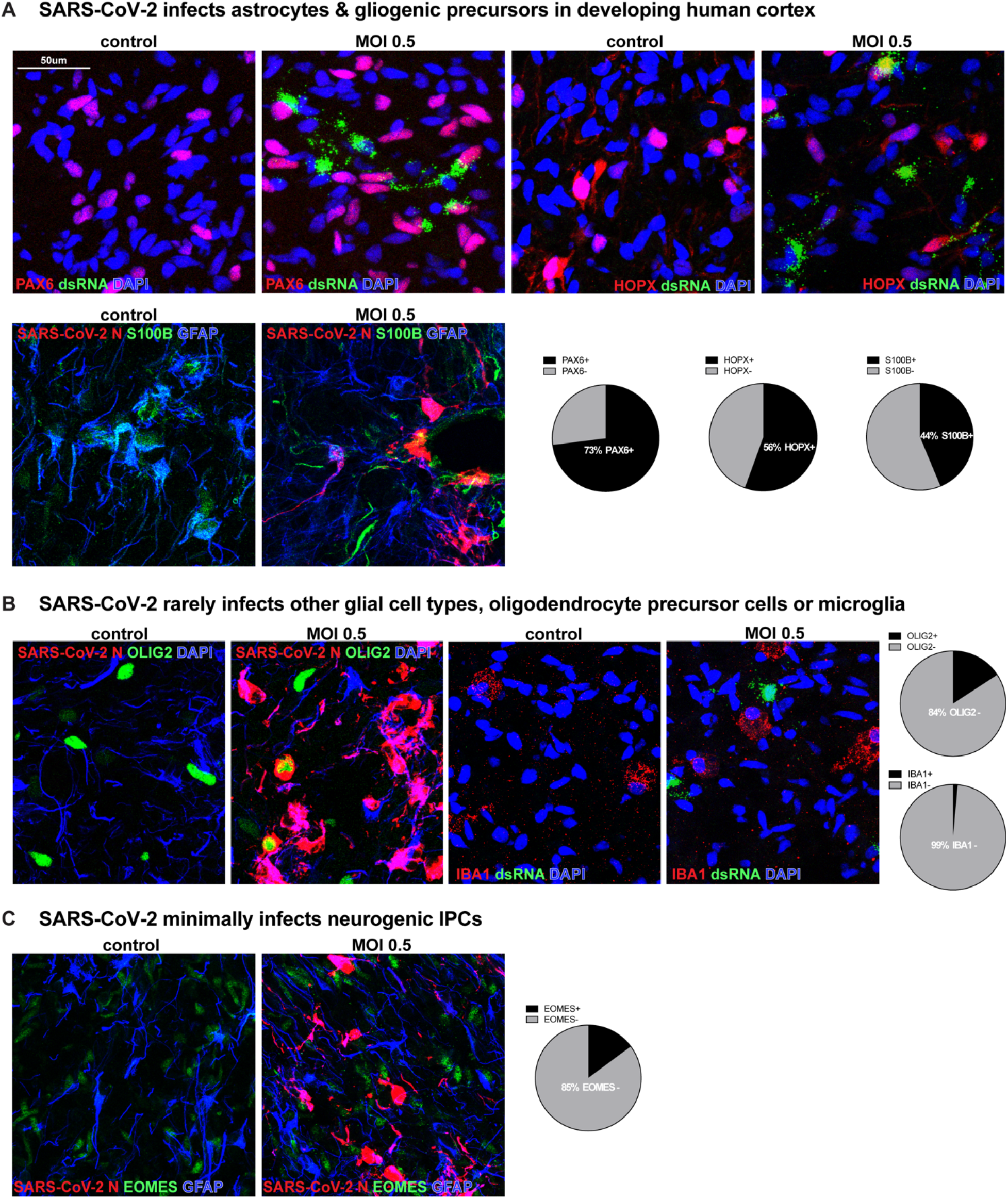
SARS-CoV-2 modestly infects other neural cell types. **A)** Mature and precursor astroglial cells indicate high infection where 74% express PAX6, 56% express HOPX and 44% express S100B (PAX6 n=252 cells, HOPX n=405 cells, S100B n=343 across two biological samples and four technical replicates). **B)** Oligodendrocyte Precursor Cells (OPC) and Microglia have little expression compared to astrocytes (OLIG2 n=487 cells, IBA1 n=350 cells). **C)** Neurogenic intermediate progenitor cells (IPCs) also demonstrates minimal infection of excitatory neuron lineage (EOMES n=128 cells).

**S Figure 2.**
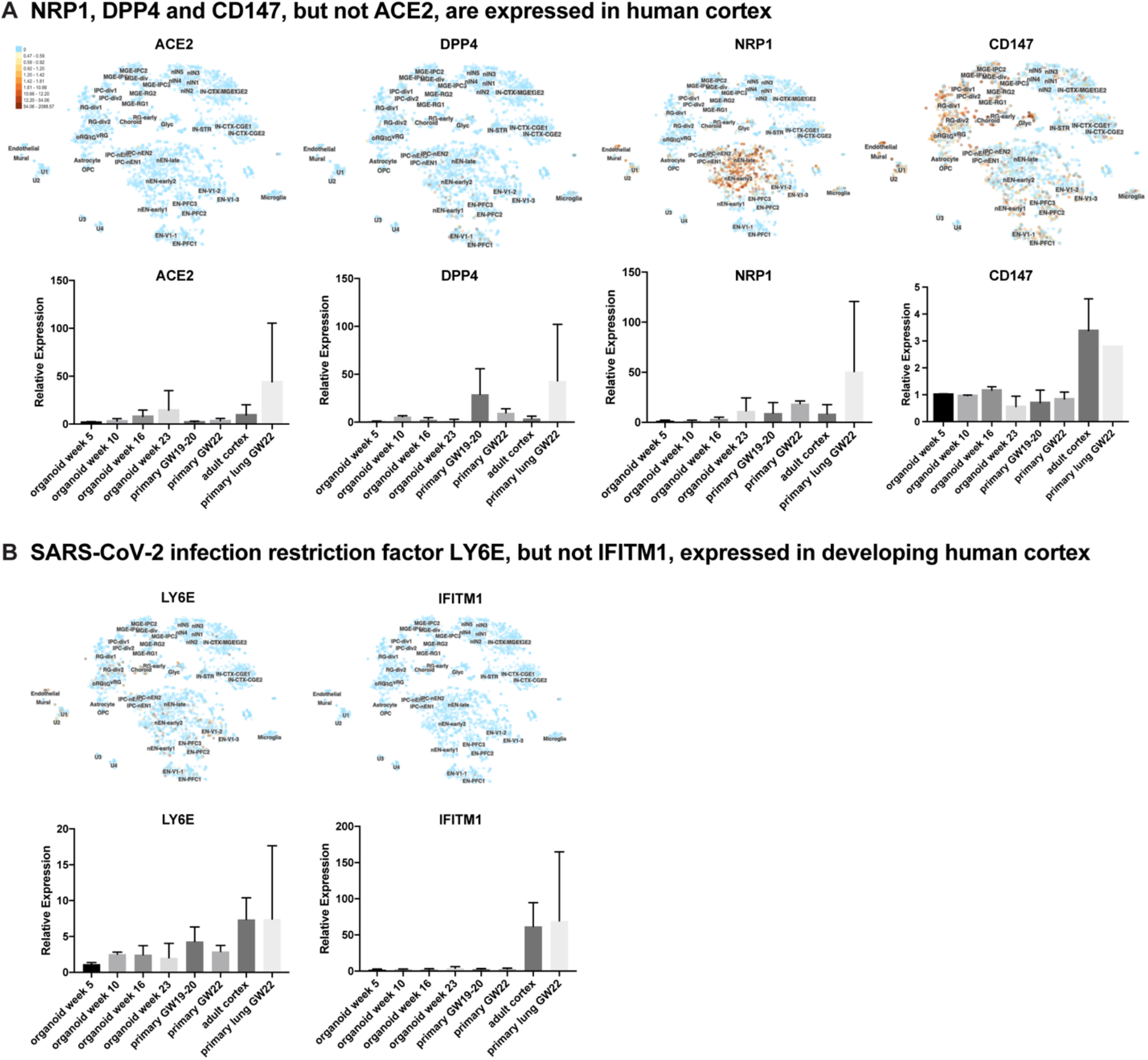
Other Coronavirus Infection Co-Factors are Expressed in Developing Human Cortex. **A)** Re-analysis of single cell RNA sequencing data from Nowakowski et al., 2017 demonstrates absent ACE2 in developing human cortex, modest DPP4, abundant CD147 across many cell types and high expression of NRP1 in excitatory neurons. Bulk RNA expression of these genes across stages of cortical organoids, developing human cortex, adult cortex and developing lung suggest there is little ACE2 RNA in the human brain. DPP4 and NRP1 are moderately expressed in bulk primary developing cortex tissue, while CD147 has highest expression in adult cortex (n>2 biological samples/sample type). **B)** Restriction factors necessary for coronavirus infection are also expressed in the developing cortex, where LY6E has broad expression and IFITM1 is only expressed in adult stages.

**S Figure 3.**
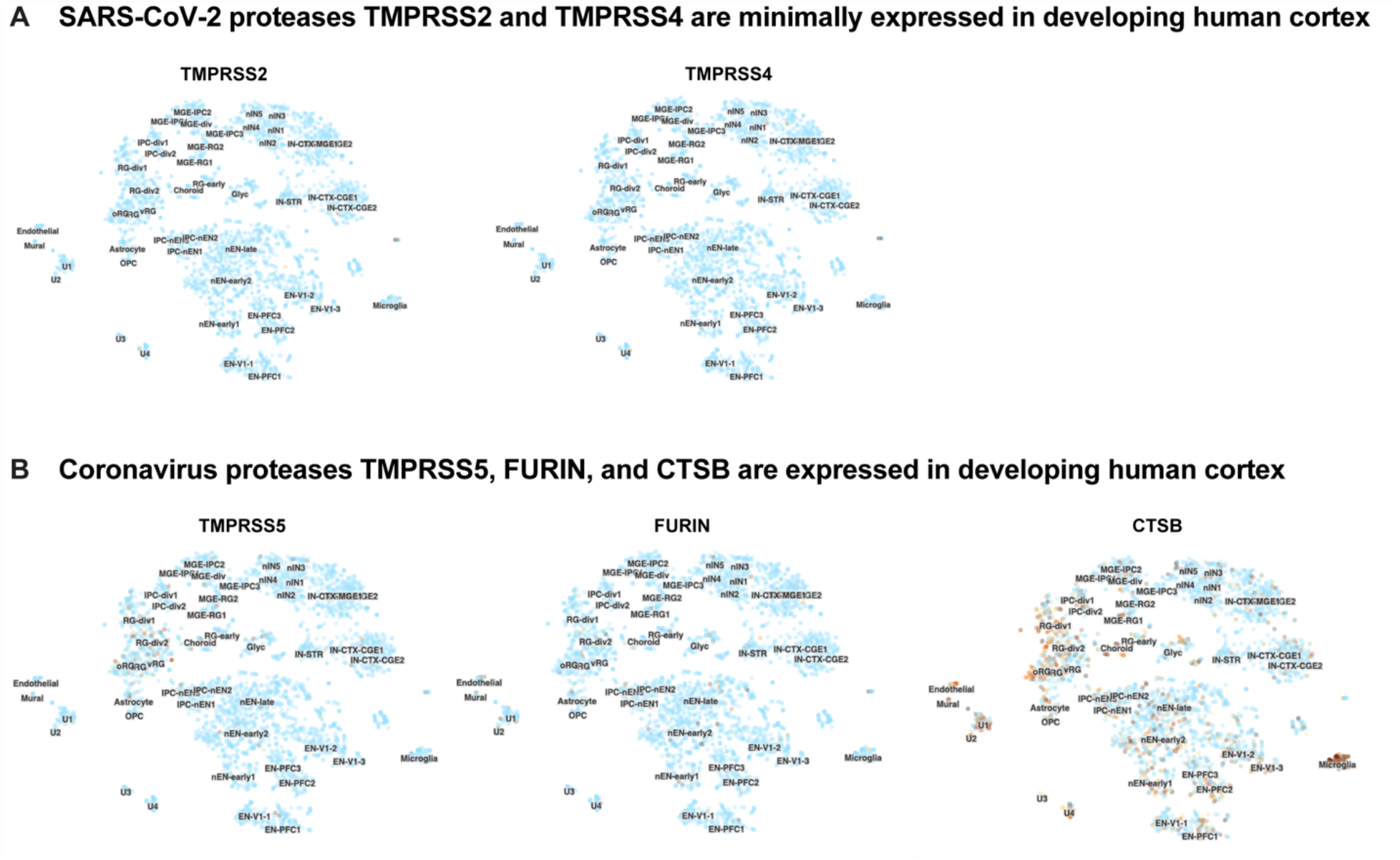
Coronavirus Protease Expression in Developing Human Cortex. **A)** Single-cell RNA sequencing data from the developing cortex demonstrates minimal expression of canonical SARS-CoV-2 proteases TMPRSS2 and TMPRSS4 in cortical cell types. **B)** Alternative coronavirus proteases, TMPRSS5, FURIN and CTSB, are differentially expressed in a variety of cell types in the developing human cortex.

## Methods

### PSC expansion culture

Human induced pluripotent stem cell lines, 13234, H28126 and H21792 (*37, 38*) and the embryonic stem cell line, H1, were expanded on growth factor-reduced Matrigel (BD)-coated six well plates. Cells were thawed in StemFlex Pro Media (Gibco) containing 10uM Rock Inhibitor Y-27632. Media was changed alternate days and lines were passaged at 70% confluency. Cells were passaged using ReLeSR™ (Stem Cell Technologies) and residual cells plated again on fresh Matrigel-coated plates. All lines in this study were between passage 25-40.

### PSC line authentication

All cell lines in this study were validated for pluripotency and karyotyped. Every 10 passages cells were tested for karyotypic abnormalities and validated for pluripotency markers Sox2, Nanog and Oct4. All cell lines tested negative for mycoplasma.

### Cortical organoid differentiation protocol

Cortical organoids were derived using a forebrain directed differentiation protocol (*37, 39*). Stem cell lines were expanded and dissociated to single cells using accutase. After dissociation, cells were reconstituted in neural induction media maintaining a density of 10,000 cells per well of 96-well V-bottom low adhesion plates. First media used is GMEM-based neural induction which includes 20% Knockout Serum Replacer (KSR), 1X non-essential amino acids, 0.11mg/mL Sodium Pyruvate, 1X Penicillin-Streptomycin, 0.1mM Beta Mercaptoethanol, 5uM SB431542 and 3uM IWR1-endo. Cells were treated with 20uM Rock inhibitor Y-27632 for the first 6 days. After 18 days, organoids were transferred to 6-well low adhesion plates and moved to an orbital shaker rotating at 90RPM and changed to DMEM/F12-based media containing 1X Glutamax, 1X N2, 1X CD Lipid Concentrate and 1X Penicillin-Streptomycin. After 35 days, organoids were moved into DMEM/F12-based media containing 10% FBS, 5ug/mL Heparin, 1X N2, 1X CD Lipid Concentrate and 0.5% Matrigel. At 70 days media was additionally supplemented with 1X B27 and Matrigel concentration increased to 1%. Throughout culture duration organoids were fed every other day. Organoids were collected for infection and RNA extraction at weeks 5, 10, 16 and 22 of differentiation.

### Organotypic slice culture

Primary cortical tissue was maintained in artificial cerebrospinal fluid (125 mm NaCl, 2.5 mm KCl, 1 mm MgCl2, 2 mm CaCl2, 1.25 mm NaH2PO4, 25 mm NaHCO3, 25 mm d-(+)-glucose) bubbled with 95% O2/5% CO2 until embedded in a 3.5% low melt agarose gel. Embedded tissue was acute sectioned at 350um using a vibratome (Leica) and plated on Millicell (Millipore) inserts in a 6 well tissue culture plate. Slices were cultured at the air liquid interface in media containing 32% Hanks BSS, 60% BME, 5% FBS, 1% glucose, 1% N2 and 1% Penicillin-Streptomycin-Glutamine. Slices were maintained for 5 days in culture at 37°C and media changed every third day.

### RNA Isolation and qPCR

Primary cortical tissue (developing and adult) and organoid samples were collected and processed for RNA extraction using QIAGEN RNeasy Plus Micro Kit. Following the QC, cDNA was prepared using SuperScript™ IV VILO™ Master Mix and qPCR performed for selected genes of interest using LightCycler^®^ 480 SYBR Green I Master Mix. Primer sequences for genes of interest are:

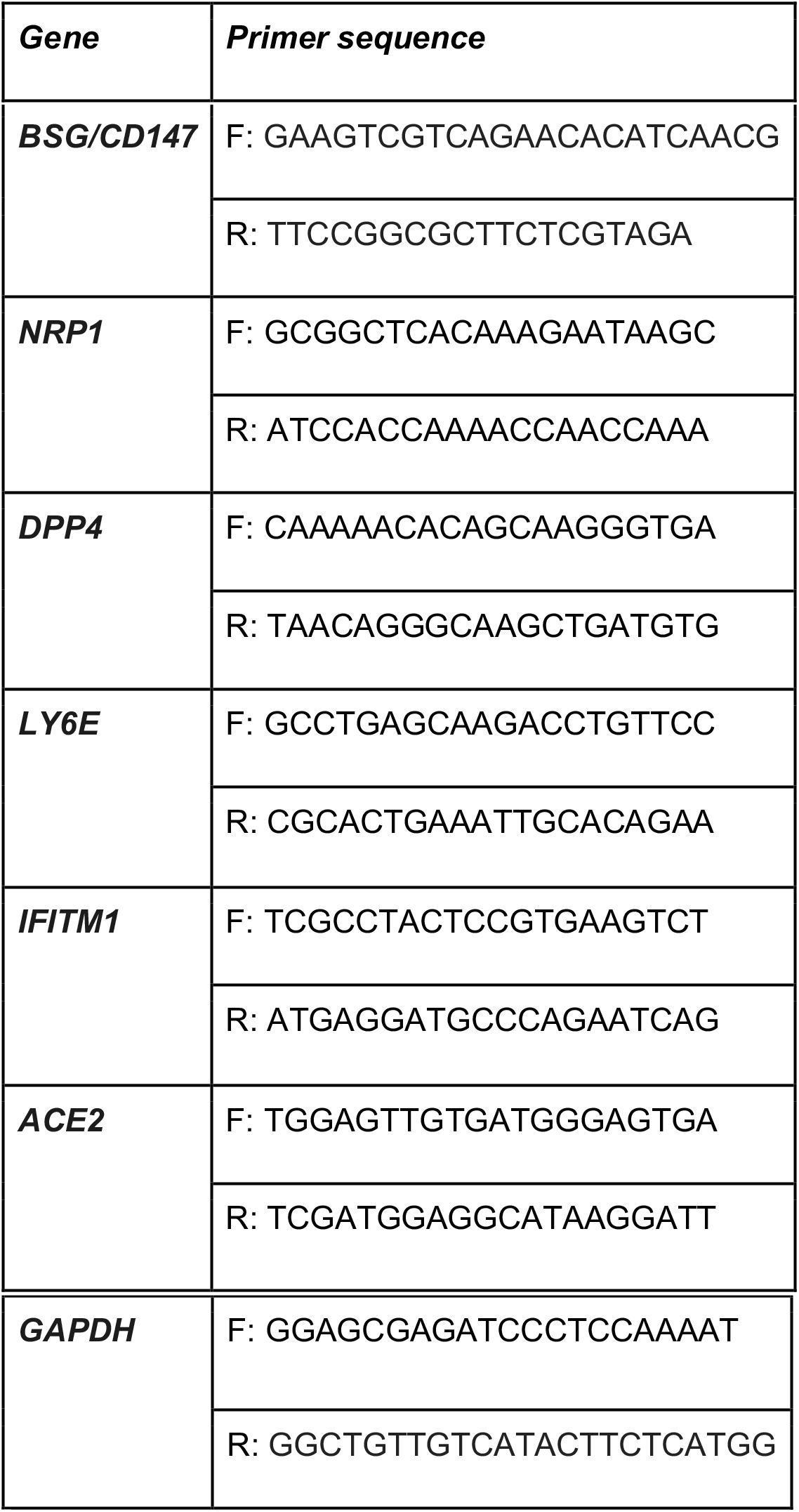

### SARS-CoV-2 Infection

The WA-1 strain (BEI resources) of SARS-CoV-2 was used for all experiments and all live virus experiments were performed in a Biosafety Level 3 lab. SARS-CoV-2 stocks were passaged in Vero cells (ATCC) and titer was determined via plaque assay on Vero cells.

SARS-CoV-2 infections were done at the indicated MOI and incubated with virus for 2 hours. After inoculation, the media was removed, cells were washed with PBS 2x, culture media was replaced, and cells were incubated at 37°C for 72 hours. Culture media was collected, samples were washed with PBS, and fixed with 4% PFA for 1 hour before removal from the BSL-3 laboratory.

For inhibitor experiments, slices were additionally cultured with 100uM Vildagliptin (Sigma, CDS022675), starting 24 hours before infection. The slices were maintained in the media plus inhibitor throughout the span of the experiment.

### Immunohistochemistry

Cortical organoids after infection were collected, fixed in 4% PFA for 1 hour, washed with 1xPBS for 2 hours and submerged in 30% sucrose in 1xPBS until saturated. Organoids were embedded in cryomolds containing 50% O.C.T. (Tissue-tek) and 50% of 30% sucrose in 1xPBS and frozen at −80C. Organoids were sectioned at 10um onto glass slides. Antigen-retrieval was performed on sections using a citrate-based antigen retrieval solution at 100x (Vector Labs) which was boiled, and tissue submerged in solution for 20mins. After antigen retrieval, slides were blocked with PBS containing 5% donkey serum, 2% gelatin and 0.1% Triton X-100 for 1 hour. Primary antibodies were incubated in blocking buffer on slides at 4°C overnight, washed with PBS containing 0.1% Triton X-100 three times and then incubated with AlexaFluor secondary (Thermo Fisher) antibodies at room temperature for 3 hours.

Organotypic slice cultures were fixed for 1 hour in 4% PFA and washed with 1xPBS 2 hours at room temperature. Slices were submerged in 30% sucrose in 1xPBS until saturated and embedded in cryomolds containing 50% O.C.T. (Tissue-tek) and 50% of 30% sucrose in 1xPBS and frozen at −80°C. Slices were sectioned at 10um onto glass slides, subjected to boiling citrate-based antigen retrieval solution (Vector Labs) for 20 min and permeabilized and blocked in blocking buffer (PBS plus 0.1% Triton X-100, 10% donkey serum, and 0.2% gelatin) for 1 h at room temperature. Primary antibodies were diluted in blocking buffer and applied to sections overnight at 4°C. Sections were washed with PBS plus 0.5% Triton X-100 and then incubated in AlexaFluor secondary antibodies (Thermo Fisher and Jackson Labs) diluted in blocking buffer at 4 °C overnight. Images were acquired on a Leica TCS SP5 X laser confocal microscope.

Primary Antibodies include: Mouse: dsRNA, clone rJ2 (Millipore, MABE1134, 1:100), Sox2 (Santa Cruz, sc-365823, 1:500), S100B (Sigma, S2532, 1:200), Ki67 (Abcam, ab156956, 1:500), CD31 (Agilent, GA61061-2, 1:100), Olig2 (Millipore, MABN50, 1:100), Rabbit: SARS-CoV-2 (Sino Biological, 40143-R001, 1:200), Pax6 (Biolegend, 901301, 1:500), Hopx (Proteintech, 11419-1-AP, 1:500), Cleaved Caspase-3 (Cell Signaling, 9661S, 1:250), Synm (Proteintech, 20735-1-AP, 1:100), Aqp4 (Proteintech, 16473-1-AP, 1:600), Egfr (Abcam, ab32077, 1:100), Dpp4 (Proteintech, 10940-1-AP, 1:50), CD147 (Invitrogen, 34-5600, 1:500), Arcn1 (Proteintech, 23843-1-AP, 1:50), Rat: Gfap (Thermofisher, 13-0300, 1:200), Laminin (Abcam, ab80580, 1:500), Nrp1 (Abcam, ab81321, 1:50), Chicken: Gfap (Abcam, ab4674, 1:500), Map2 (Abcam, ab5392, 1:200), Goat: Ace2 (R&D, AF933, 1:200), Ace2 (Thermofisher, MA5-32307, 1:200), Iba1 (Abcam, ab48004, 1:500), Pdgfrb (R&D, AF385, 1:100), Sheep: Eomes (R&D, AF6166, 1:200), Guinea pig: NeuN (Millipore, ABN90, 1:500), Sheep: CD34 (R&D, AF7227, 1:200).

